# Phylogenetic ANOVA: The Expression Variance and Evolution (EVE) model for quantitative trait evolution

**DOI:** 10.1101/004374

**Authors:** Rori V. Rohlfs, Rasmus Nielsen

## Abstract

A number of methods have been developed for modeling the evolution of a quantitative trait on a phylogeny. These methods have received renewed interest in the context of genome-wide studies of gene expression, in which the expression levels of many genes can be modeled as quantitative traits. We here develop a new method for joint analyses of quantitative traits within and between-species, the Expression Variance and Evolution (EVE) model. The model parameterizes the ratio of population to evolutionary expression variance, facilitating a wide variety of analyses, including a test for lineage-specific shifts in expression level, and a phylogenetic ANOVA that can detect genes with increased or decreased ratios of expression divergence to diversity, analogous to the famous HKA test used to detect selection at the DNA level. We use simulations to explore the properties of these tests under a variety of circumstances and show that the phylogenetic ANOVA is more accurate than the standard ANOVA (no accounting for phylogeny) sometimes used in transcriptomics. We then apply the EVE model to a mammalian phylogeny of 15 species typed for expression levels in liver tissue. We identify genes with high expression divergence between-species as candidates for expression level adaptation, and genes with high expression diversity within-species as candidates for expression level conservation and/or plasticity. Using the test for lineage-specific expression shifts, we identify several candidate genes for expression level adaptation on the catarrhine and human lineages, including genes putatively related to dietary changes in humans. We compare these results to those reported previously using a model which ignores expression variance within-species, uncovering important differences in performance. We demonstrate the necessity for a phylogenetic model in comparative expression studies and show the utility of the EVE model to detect expression divergence, diversity, and branch-specific shifts.

---

Quantitative phylogenetic methods account for non-independence relationships between species using several approaches such as independent contrasts (Felsenstein 1985) and generalized least squares (Grafen 1989; Martins and Hansen 1997; Rohlf 2001). These methods have provided frameworks for a variety of phylogenetic approaches which consider variance within species (for a review, see Garamszegi 2014). For instance, the phylogenetic mixed model considers both gradual evolutionary drift and within-species variance (Lynch 1991; Housworth et al. 2004). Another approach transforms comparative quantitative data to account for phylogeny before performing ANOVA (Butler et al. 2000). Still other methods compare ANOVA results based on raw phylogenetic data to those based on data simulated under a phylogenetic model to create an appropriate null distribution (Garland et al. 1993; Harmon et al. 2008; Revell 2012). Sophisticated extensions of quantitative evolutionary models allow evolutionary scenarios including varying rates of phenotypic evolution (Pagel 1999; O’Meara et al. 2006). These quantitative trait evolution methods have been used effectively for a variety of phenotypic, particularly morphological, traits.

The emergence of transcriptome-wide comparative gene expression studies including multiple individuals per species (Kalinka et al. 2010; Brawand et al. 2011; Perry et al. 2012; Necsulea et al. 2014) has presented a new challenge to quantitative evolutionary methodology. Like traditional morphological traits, expression levels can be considered a quantitative trait that evolves over a phylogeny. Expression levels are particularly interesting as relatively malleable basic genetic traits, creating a convenient point of intervention for adaptation (Whitehead and Crawford 2006; Gilad et al. 2006a; Fraser 2011). By examining comparative expression levels, we can identify fundamental changes that underlie adaptation to environmental factors. This invites quantitative genetic investigation of evolutionary modality (drift, stabilizing selection, adaptive shift, etc.). In addition to a clear genetic basis, expression levels have strong environmental components (Idaghdour et al. 2010; Pickrell et al. 2010). Changes in expression level may reflect genetic adaptation fixed within individuals, or plastic (rapidly changeable) response to environmental variables. This plasticity allows examination of the relationship between expression plasticity and adaptability. Finally, the large numbers of measurements across genes in transcriptome-wide expression studies present new analytical opportunities.

Despite the extensive literature of quantitative phylogenetic methods, many early large-scale comparative expression analyses used traditional ANOVA to detect genes with unusually high expression divergence between-species, given the expression variance within-species (Nuzhdin et al. 2004; Gilad et al. 2006b; Khaitovich et al. 2006; Whitehead and Crawford 2006). These analyses typically assume independence between species. While technically untrue, this assumption has no impact for phylogenies of two species and may have limited impact for the small numbers of species analyzed. However, as more species are considered in recent studies, the difference in shared evolutionary history between closely and distantly related species increases, and a complex covariance structure emerges. In current comparative expression datasets across larger phylogenies, the assumption of species independence does not hold, necessitating more sophisticated methods taking into account evolutionary relationships (Felsenstein 1985).

More recent comparative expression studies have employed classical quantitative trait evolutionary models, particularly the model of constrained trait evolution proposed by Hansen (1997) and expanded in later work (Butler and King 2004; Hansen et al. 2008). This flexible model has been applied to describe the evolution of gene expression under neutral expression level diffusion, constrained diffusion (expected under stabilizing selection), and species-specific expression level shifts (Bedford and Hartl 2009). These models are used to calculate the expected species average expression levels and expression covariance between species under a particular evolutionary scenario. Likelihood ratio tests can then be formulated to distinguish unconstrained random trait evolution, constrained or stabilized trait evolution, and branch-specific shifts in trait evolution, as has been successfully analyzed in a number of datasets (Bedford and Hartl 2009; Kalinka et al. 2010; Perry et al. 2012; Schraiber et al. 2013). However, these methods are limited by their inability to model non-phylogenetic variance (Oakley et al. 2005) and are not designed to investigate evolutionary expression variation in relation to expression variance within-species.

A number of augmentations to these models allow within-species variance as an error term (Martins and Hansen 1997; Lynch 1991; Gu 2004; Ives et al. 2007; Felsenstein 2008; Hansen and Bartoszek 2012; Rohlfs et al. 2014). Several models of phenotypic drift parameterize within-species variance (Lynch 1991; Housworth et al. 2004; Felsenstein 2008), while other analyses show how this substantially improves ancestral state estimation (Martins and Lamont 1998; Ives et al. 2007) and evolutionary inference (Harmon and Losos 2005; Ives et al. 2007; Revell et al. 2008). Within-species variance has additionally been parameterized in an evolutionary model allowing for constrained trait evolution (Rohlfs et al. 2014).

We build upon these models to create the unified Expression Variance and Evolution (EVE) model, describing both phylogenetic expression level evolution between species and expression level variance within-species. Expression levels vary among individuals in a population or a species. This expression level variance is caused by genetic and environmental differences among individuals. It may be low if the gene has an important function, is expressed constitutively, and does not respond to environmental changes. Such genes might be genes involved in important cellular functions such as cell cycle control. Genes that have high expression level variance are genes that either harbor segregating adaptive variation affecting expression levels, or more likely, respond to various environmental cues. Such genes might, for example, include genes involved in immunity and defense against pathogens. Our method allows for expression level evolution under neutrality or selective constraint with a flexible model (Hansen 1997; Butler and King 2004; Hansen et al. 2008), while adding in within-species variance (as was previously done under drift (Lynch 1991; Housworth et al. 2004; Felsenstein 2008)). The EVE model re-parameterizes a previous model which allows within-species variance simply as an error term (Rohlfs et al. 2014). By contrast, in the EVE model, we parameterize the ratio of expression variance within-species to evolutionary variance between-species, facilitating rigorous novel analyses directly aimed at this ratio. This can be considered a phylogenetic analogy to test for drift via ratios of between- to within-population variance Lande (1979); Ackermann and Cheverud (2002); Marroig and Cheverud (2004). We develop this phylogenetic framework with genome-wide expression data in mind, exploiting the large number of expression measurements over the same individuals. Yet, the EVE model could be used for any set of quantitative traits, including morphological traits.

The EVE model enables an expression analogy to classic genetic neutrality tests considering polymorphism and diversity, namely, the HKA test (Hudson et al. 1987). In this test, the ratio of polymorphism within-species to divergence between-species is compared among different genes in the genome. Under neutrality, this ratio should be the same (in expectation) for all genes in the genome. However, for genes affected by selection, the number of polymorphic sites within-species may be increased or decreased relative to the number of fixed differences between-species, depending on the directionality and modality of selection (see e.g., Nielsen 2005).

Analogously, in our model, we parameterize the ratio of within-species expression variance to between-species expression evolutionary variance using a parameter β defined over the phylogeny. This parameter represents the ratio of within-to between-species variance, which should be approximately constant for a given phylogeny over different genes if only constant stabilizing selection (or no selection) is acting on the trait (Lande 1976). We can now construct likelihood ratio tests aimed at detecting if *β* varies among genes. Let *G* = *g*_1_, *g*_2_, …, *g*_*k*_ be the set of all *k* genes for which expression values have been obtained, and let the value of *β* for gene *i ∈ G* be *β*_*i*_. To test if *β*_*i*_ is elevated compared to the rest of the genes, we then calculate the likelihood under the null hypothesis of a constant value of *β* among genes, i.e. *β*_*i*_ = *β*_*shared*_ for all genes *i* ∈ *G*. We compare it to the alternative hypothesis of *β*_*i*_ ≠ *β*_*shared-i*_, where *β*_*shared-i*_ is a value of *β* shared for all genes in *G* except *g*_*i*_. The resulting likelihood ratio test statistic, formed in the usual fashion, by comparing the log likelihood maximized under the union of the null and the alternative hypothesis, to the log likelihood maximized under the null hypothesis, is then chi-square distributed with one degree of freedom under standard regularity conditions.

As a practical matter, we assume that the value of *β* estimated for *β*_*shared*_ is approximately the same as the value of *β* estimated for *β*_*shared-i*_ for any *i*. This assumption is reasonable when there are many genes and the estimate of *β*_*shared*_ is not dominated by any particular gene. Using this assumption leads to considerable reductions in computational time. In the following, we will therefore in the notation not distinguish between *β*_*shared*_ and *β*_*shared-i*_.

If the null hypothesis is rejected because *β*_*i*_ is significantly larger than *β_shared_*, expression divergence between-species is elevated in gene *i* relative to the level of within-species variance. This would suggest that gene *i* may be subject to species or branch-specific directional selection on expression level. Genes with an unusually low ratio (*β*_*i*_ < *β*_*shared*_) show proportionally high expression diversity within-species, suggesting conservation of species average expression levels, with expression variation in response to either environmental factors or diversifying selection within species. This test can also be thought of as an alternative phylogenetic ANOVA test as it is essentially an analysis of expression variance within-versus between-species, accounting for varying evolutionary relationships between species. In statistical terms, the analogy is to a one way ANOVA where species define the discriminating factor and the test determines if species share the same mean, but where evolutionary dependencies between species are accounted for.

Since phylogenetic information is included in the EVE model itself, a wide variety of evolutionary scenarios may be specified by selectively constraining parameters, improving flexibility to test different comparative hypotheses. For example, we can test for unusual species or lineage-specific expression variance, as may be observed under recent relaxation or increases of constraint on expression level, diversifying selection on expression level, or under extreme branch-specific demographic processes. Other tests may be constructed to test for differing expression diversity for groups of individuals within each species, for instance, evolutionarily conserved age or sex-specific expression variance. All of these tests could be performed on a particular gene of interest or on a class of genes of interest, for example, a list of candidate genes could be queried for increased expression diversity in older individuals. In addition to these novel tests, the EVE model can be used for the same tests as other expression evolution models which discount within-species variance. In particular, the EVE model can test for lineage-specific shifts in constrained expression level, while taking into account within-species variance.

Here, we explore the performance of two EVE model tests: the test for unusual expression divergence or diversity and the test for lineage-specific expression level shifts. We use simulations to describe these tests and formulate expectations under the null hypotheses. We then apply the tests to a previously published expression dataset of 15 mammals. We identify a number of genes with high expression level divergence between-species as candidates for expression level adaptation to species-specific factors, and genes with high expression level diversity within-species as candidates for environmentally responsive gene expression (plasticity). Using the test for lineage-specific expression shifts, we identify several strong candidate genes for branch-specific expression adaptation on the catarrhine and human lineages.

We compare our results to those obtained using the species mean model described by Bedford and Hartl (2008) and recently used in a number of studies (Bedford and Hartl 2009; Kalinka et al. 2010; Perry et al. 2012). The species mean model considers the evolution of the mean expression level for each species, rather than within-species variance. This model can describe trait evolution without constraint, with constraint, or with a branch-specific adaptive shift in response to an environmental factor. By comparing the likelihood of observed data under different parametric limits, the species mean model can be used to identify genes subject to different evolutionary schemes. We find important differences between our results and those obtained using the species mean method, especially for analyses of species-specific expression shifts (Perry et al. 2012).

## Methods

### The EVE Model for Gene Expression Evolution and Population Variance

The evolution of quantitative traits by diffusion and constrained or stabilized diffusion has been modeled using an Ornstein-Uhlenbeck (OU) process, which can be thought of as a random walk with a pull towards an optimal value (Lande 1976; Hansen 1997; Butler and King 2004; Hansen et al. 2008; Bedford and Hartl 2009; Kalinka et al. 2010). In an OU model of stabilizing selection on gene expression level, the parameter *θ*_*i*_ can be thought of as the optimal expression level for gene *i*, 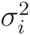 the diffusion acting on that expression level, and *α*_*i*_ the rate of adaptation for that expression level (Hansen 1997; Butler and King 2004; Hansen et al. 2008; Hansen 2012). Over evolutionary time, the stationary variance of species mean expression levels for gene *i* will be 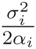, which we refer to as the evolutionary variance.

More recently, several Brownian motion and OU-based models have been augmented to include within-species population level variance (Felsenstein 2008; Lynch 1991; Hansen and Bartoszek 2012; Rohlfs et al. 2014). Accounting for population variance is crucial to distinguish evolutionary modalities (Rohlfs et al. 2014).

The model we describe builds on these OU models for quantitative trait evolution with the additional parameter *β* which describes the ratio of population to evolutionary expression level variance. Within species *j* the expression level of any individual *k* is distributed as 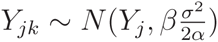, where *Y*_*j*_ is the species mean expression level determined by the OU process. We call this the EVE model, which describes a linear relationship between population and evolutionary expression level variance.

In his classic paper, Lande (1976) showed that under an OU model of stabilizing selection, a linear relationship arises between a quantitative trait’s evolutionary variance and population variance within-species. Additionally, the Poisson nature of RNA-Seq and gene expression itself means that both evolutionary and population expression variance increase with expression mean. With that in mind, our model assumes a linear relationship between evolutionary and population expression variance. That assumption is reflected in the data, which shows a linear relationship between estimated evolutionary expression level variance 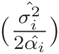 and estimated population expression level variance 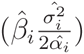 (Figure 1).

**Figure 1:** The maximum likelihood estimated per-gene evolutionary variance 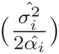 and population variance 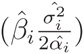 are plotted against each other. The linear regression line is shown.

The slope of this linear relationship (parameterized by *β*) should be consistent across genes which have undergone the same evolutionary and demographic processes under stabilizing selection. However, in a gene, *i*, which has experienced directional selection on expression level, *β*_*i*_ would be lower as compared to other genes in the same individuals. The directional selection would drive increased expression divergence between-species, while maintaining low expression variance within-species. Similarly, a gene with plastic expression may have more variation within-species than between as compared to other genes, raising the value of *β*_*i*_. High *β*_*i*_ could alternatively be explained by diversifying selection on expression level. Since expression levels are quite plastic, this explanation seems less plausible without other corroborating information. In this manuscript, since the samples we consider are opportunistically harvested, presumably under quite varying environmental conditions, we focus on the environmental plasticity hypothesis in the interpretation of our results.

### Likelihood Calculations Under the EVE Model

The EVE model is similar to other OU-process-based phylogenetic models (Butler and King 2004; Bedford and Hartl 2009), with the addition of within-species expression variance in terms of the evolutionary variance. As such, under the EVE model expression levels across individuals and species, given a fixed phylogeny, follow a multivariate normal distribution identical to those under species means models at the species level as

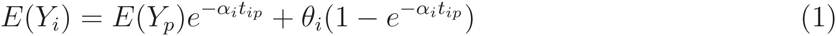

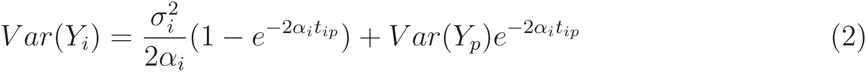

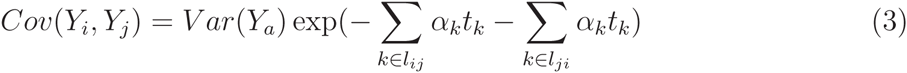

where *Y*_*i*_ is the expression level in species *i*; *Y*_*p*_ is the species mean expression at the parental node *p* of species *i*; *θ*_*i*_, 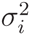, and *α*_*i*_ are the parameter values on the branch leading to node *i*; *t*_*ip*_ is the length of the branch between *i* and *p*; *Y*_*a*_ is the expression level at the most recent common ancestor of species *i* and *j*; and *l*_*ij*_ is the set of nodes in the lineage of *Y*_*i*_ not in the lineage of *Y*_*j*_ (Rohlfs et al. 2014).

This multivariate normal distribution describing the species-level expression is augmented in the EVE model to include individuals within species, so for an individual *k* in species *i*, 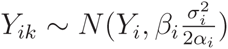. In this way, the within-species expression variance parameter described by Rohlfs et al. (Rohlfs et al. 2014) τ^2^ is re-parameterized as 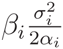.

The entire multivariate normal distribution can be described as

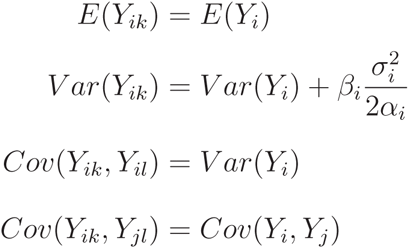

based on equations 1, 2, and 3, where *i* ≠ *j* and *k* ≠ *l*. With the distribution of expression levels under a particular set of parameters defined according to this multivariate normal, the likelihood of the data under the model is simply the probability density. Notice that sampling and experimental variance is accounted for (and confounded) in the parameters governing the distribution of *Y*_*ik*_|*Y*_*i*_.

### Maximum Likelihood Procedures

For the test for individual gene departures from *β*_*shared*_, under the null hypothesis each gene *i* is governed by parameters *θ*_*i*_, 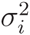, and *α*_*i*_, reflecting the evolutionary process of each gene based on its degree of expression diffusion and constraint. The population expression variance in all *n* genes is controlled by the single parameter *β*_*shared*_. To more computationally efficiently maximize the likelihood over these 3*n* + 1 parameters, we use a nested structure with Brent’s method (Brent 1973) in the outer loop to maximize over the single parameter *β*_*shared*_, and the BFGS algorithm (Broyden 1970; Fletcher 1970; Goldfarb 1970; Shanno 1970) in the inner loop to optimize over *θ*_*i*_, 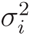, and *α*_*i*_ for each gene. Under the alternative hypothesis, the likelihood of each gene *i* is maximized using the BFGS algorithm over *θ*_*i*_, 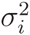, *α*_*i*_, and *β*_*i*_. To compute the likelihood ratio, the likelihoods of each individual gene *i* are computed under *H*_0_ : *β*_*i*_ = *β*_*shared*_ and *H*_*a*_ : *β*_*i*_ ≠ *β*_*shared*_, where *β*_*shared*_ considers all of the genes considered. Note that this experimental set up allows better computational efficiency, but relies on *β*_*shared*_ over all the genes approximating *β*_*shared*_ over all the genes excluding gene *i* for large numbers of genes.

In the likelihood maximization under the null hypothesis, likelihoods across genes are assumed to be independent so that for a particular value of *β*_*shared*_, the likelihood of a set of genes is simply the product of the likelihoods of each gene. While this assumption is currently typical in this sort of analysis, it leaves something to be desired since the evolution of expression levels of inter-related genes are not independent, and nor are the particular expression levels measured in an individual which may be responding to the environment of that individual. A more rigorous approach would take into account complex correlation structures across genes, as has been outlined for some evolutionary models (Lande and Arnold 1983; Felsenstein 1985, 1988; Lynch 1991). Unfortunately, because of the combinatorial problem of investigating a very large set of possible correlation structures, a full likelihood approach that estimates the correlation structure directly for thousands of genes is not computationally tractable and possibly may not be based on identifiable models. Instead we use the independence model as an approximation. If expression patterns are correlated among genes, we can consider this procedure to be a composite likelihood method (Larribe and Fearnhead 2011) since the estimating function is formed by taking the product of functions that individually are valid likelihood functions, but the total product is not necessarily a valid likelihood function. In the case of severe dependence between genes, estimates of *β*_*shared*_ will tend towards the value for correlated genes, leading to over-identification of genes with *β*_*i*_ different from the correlated genes.

For the test of branch-specific expression shift for a particular gene *i*, under the null hypothesis the likelihood of each gene *i* is maximized over *θ*_*i*_, 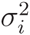, *α*_*i*_, and *β*_*i*_. Under the alternative hypothesis the likelihood of each gene *i* is maximized with an additional *θ* parameter 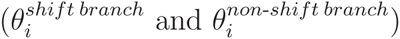 to allow for the expression shift.

### Testing for deviations from a constant expression divergence/diversity ratio

The EVE model can, as previously mentioned, be used to test for deviations from a constant ratio of expression to divergence ratio among genes, analogous to the HKA test often applied to test for selection at the DNA level. Specifically, a likelihood ratio can be formed by comparing the likelihood under a null model where *β* for all genes equals *β*_*shared*_ (H_0_ : *β*_*i*_ = *β*_*shared*_) to the likelihood under the alternative model where *β*_*i*_ is a free parameter (*H*_*a*_ : *β*_*i*_ ≠ *β*_*shared*_). If the null hypothesis is rejected in a likelihood ratio test, we can conclude that *β*_*i*_ for a particular gene varies significantly from *β*_*shared*_ across the genes. A gene where *β*_*i*_ < *β*_*shared*_ has high expression variance between-species as compared to within, or high expression divergence. A gene where *β*_*i*_ > *β*_*shared*_ has high expression variance within-species as compared to between, or high expression diversity.

An implementation of the EVE model is available in the supplement of this paper.

### Mammalian expression data and phylogeny

We applied the EVE model to analyze a comparative expression dataset over 15 mammalian species with four individuals per species (except for armadillos with two individuals) which is described in full in Perry et al. (2012). Of the 15 species typed, five are anthropoids (common marmoset (mr), vervet (ve), rhesus macaque (mc), chimpanzee (ch), human (hu)), five are lemurs (aye-aye (ay), Coquerel’s sifaka (sf), black and white ruffed lemur (bw), mongoose lemur (mn), and crowned lemur (cr)), and the remaining five are more distantly related mammals (slow loris (sl), northern treeshrew (ts), house mouse (ms), nine-banded armadillo (ar), and gray short-tailed opossum (op)). Since many of these species are endangered and protected, most samples were collected opportunistically within four hours of death. Liver tissue from each individual was typed using RNA-Seq and transcriptomes were assembled with a robust de novo technique that was verified on species with reference genomes available (Perry et al. 2012). Expression levels were normalized based on each individual, transcript length, GC content, and species (Bullard et al. 2010; Pickrell et al. 2010; Perry et al. 2012), as is appropriate for comparative analysis so that genes are considered equitably in relation to each other (Dunn et al. 2013). Here, we consider a subset of 675 genes with no missing data across all species and individuals.

### Simulated data

#### Comparing EVE and ANOVA

We performed a simulation study to compare the power of the EVE method and traditional ANOVA to detect expression divergence between-species. Expression was simulated for 100 genes on the phylogeny and number of individuals observed experimentally, using the parameter values σ^2^ = 5, *α*= 3.0, *β*= 6, and *θ* = 100 with a total tree height of 0.08. However, one of the simulated genes was subject to a branch-specific expression shift on either the opossum, human, or anthropoid branches. These simulations were performed for varying strengths of branch-specific shifts and for each shifts on each of the three branches considered with 100 simulations in each set of conditions. For the opossum branch shift, differences in optimal expression levels (Δ *θ*) ranged from 0 to 19; for the human branch shift, values ranged from 0 to 950; and for the anthropoid branch shift, values ranged from 0 to 57. These parameter values describe relatively weak stabilizing selection with drastic branch-specific optimum shifts. The varying optimum shift values were chosen to achieve similar absolute expression level changes across the three trials with shifts on differently-lengthed branches.

#### Null distribution of LRT_*βi* = *βshared*_

We performed a second simulation study to explore the null distribution of the test statistic for unusual expression divergence or diversity (*LRT*_*βi = βshared*_). Since the alternative hypothesis has one additional degree of freedom as compared to the null hypothesis, the asymptotic distribution for the LR test statistic under the null hypothesis is chi squared with one degree of freedom 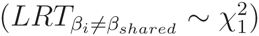.

However, smaller phylogenies may not be large enough for the asymptotic distribution to apply, as has been observed in other comparative methods (Boettiger et al. 2012; Beaulieu et al. 2012).

For our simulations exploring the null distribution of *LRT*_*β_i_*= *βshared*_, we consider a phylogeny identical to that from the mammalian dataset from Perry et al. (Perry et al. 2012), calling that “1x tree” or *t*^1^. We additionally consider a “2x tree” or *t*^2^ which is constructed with two copies of *t*^1^ as (*t*^1^, *t*^1^) with the connecting branches the length of *t*^1^ itself. Similarly, we consider a “3x tree” or *t*^3^ as (*t*^2^, *t*^1^) with the branch to *t*^2^ the length of *t*^1^ and the branch to *t*^2^ twice the length of *t*^1^, and a “4x tree” or *t*^4^ as (*t*^2^, *t*^2^) with the connecting branches the length of *t*^1^ (Supplementary Figure ??).

We performed additional simulations based on a pectinate topography over different number of species with the same internal branch lengths (for example, Supplementary Figure ??) and a single set of parameters taken from the median parameter estimates from the experimental analysis (*θ* = 0.57, σ^2^ = 2.66, α= 19.05, and *β* = 0.39).

## Results

### Comparison to traditional ANOVA

Both the traditional ANOVA and the EVE ‘phylogenetic ANOVA’ tests were performed on simulated data (described above), the later leveraging variance information over genes in addition to phylogenetic information. Figure 2 compares the power of the ‘phylogenetic ANOVA’ and traditional ANOVA. Without taking phylogeny into account, the traditional ANOVA interprets species differences attributable to drift as due to divergence, leading to uncontrolled false positive rates (Figure 2 at average expression difference of zero). The ‘phylogenetic ANOVA’ gains power for genes with moderate expression shifts by considering these shifts in the context of the phylogeny. Among the simulations with shifts on different branches, the EVE method has more power to detect shifts in the opossum lineage than the human lineage, analogous to power differences across branch lengths in sequence-based tests for divergence (Yang and dos Reis 2011). With both methods, the shift on the anthropoid lineage which includes five species is more easily detected than the single species shifts.

**Figure 2:** Power is shown as a function of average expression difference between the species on the shifted branch and the rest of the phylogeny. Power is shown for traditional ANOVA (crosses) and the EVE method ‘phylogenetic ANOVA’ (triangles) for shifts on the (a) opossum, (b) human, and (c) anthropoid branches.

### Determining significant deviations of expression divergence/diversity ratio

#### Test expectation under the null hypothesis

At the asymptotic limit, the likelihood ratio test statistic for testing *H*_*0*_ : *β*_*i*_ = *β*_*shared*_ versus *H*_*A*_ : *β*_*i*_ ≠ *β*_*shared*_, *LRT* _*β*_*i*_≠ *β_shared_*_, is 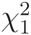 distributed under the null hypothesis. However, when applied to small phylogenies, the distribution of *LRT*_*β*_*i*_≠*β*_*shared*__ may not be near the asymptotic limit, and may deviate from a 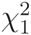 (e.g., Boettiger et al. 2012) (see Supplementary Materials). To explore the null distribution of *LRT*_*β*_*i*_≠*β*_*shared*__ over different parameter values and phylogeny sizes, we simulated data under the null hypothesis of *H*_0_ : *β*_*i*_ = *β*_*shared*_ for four sets of parameter values (Supplementary Table 1) based on the median maximum likelihood estimates from the experimental data, under four tree sizes based on the mammalian phylogeny that we subsequently will analyze (Supplementary Figure 1 and Supplementary Materials).

While the null distribution resembles the asymptotically expected 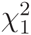 for a phylogeny like the one analyzed here, we observe some minor deviations (Supplementary Figure 2). However, as the size of the phylogeny considered increases, the null distribution approaches a 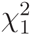, though it converges more slowly under some parameter values. As in previous studies examining parameter estimates over phylogeny size (Boettiger et al. 2012), we see that the parameter estimates improve with phylogeny height and number of tips, though some are more easily estimable than others (Supplementary Figures 5-10). Yet, note that for the set of expression values simulated under a low value (set 3), the evolutionary variance is very high and is not saturated in the phylogeny lengths explored here. In this case, the phylogenies with longer branches investigated allow more time for expression levels to vary more widely, making parameter and likelihood estimation less accurate. This is a case where the null distribution of *LRT*_*β*_*i*_ = *β*_*shared*__ is far from the asymptotic expectation.

We performed further simulations based on a pectinate phylogeny for different numbers of species (Supplementary Figures 3, 11). Again, we see that as the phylogeny size increases, the simulated null distribution more closely matches the asymptotic expectation. It is important to note that the null distribution under a pectinate topology more quickly approaches 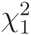 than the other topology because there are more varying branch lengths between species in a pectinate phylogeny. Trait evolution methods are powered by multiple varying branch length differences between species, making a pectinate phylogeny the most informative.

### Parametric bootstrap approach for the null distribution

To account for deviations from the asymptotically expected null distributions of *LRT*_*β*_*i*_≠*β*_*shared*__, we follow the suggestion of Boettiger et al. (2012) and use a parametric bootstrap. That is, for a particular gene, we simulate expression profiles based on the maximum likelihood parameter estimates under the null hypothesis. These simulated expression profiles are then tested for deviation from the null hypothesis to determine the parametric bootstrapped null distribution of *LRT*_*β*_*i*_≠*β*_*shared*__, to which the experimental result can be compared.

We performed a parametric bootstrap analysis with 100 simulations for each of the genes simulated under the null hypothesis described above. For each gene, we compared the original test statistic (*LRT*_*β*_*i*_≠*β*_*shared*__) to the distribution created by these additional simulations to determine the parametric bootstrapped *p*-value. The resulting bootstrapped *p*-values are approximately uniformly distributed between 0 and 1 (Supplementary Figure 13) as expected. Note that these bootstrapped *p*-values describe the departure from the null for each gene individually; a correction for multiple tests must be included when considering *p*-values across genes. Further, note that the bootstrap approach assumes independence between genes, which, while statistically convenient, could cause inaccuracy when expression is highly correlated between genes. Generally the parametric bootstrap approach is most effective for accurate parameter estimates; in the presence of biased estimates and a dependence of the distribution of the likelihood ratio test statistics on parameter values, the parametric bootstrap approach can be biased. It is therefore worthwhile to test the parametric bootstrap before interpreting results based on it.

### Expression Divergence and Diversity in Mammals

#### Assessing expression divergence and diversity

We applied the test of constant expression divergence to diversity ratio to each gene in the mammalian dataset. The resulting empirical *LRT*_*β*_*i*_≠*β*_*shared*__ values increase with departure from 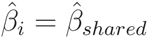 (Figure 3). We see much higher values of *LRT*_*β*_*i*_≠*β*_*shared*__ for low 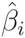 than high 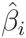. This is partially explained by error in *β*_*i*_ estimates, especially for higher values (Supplementary Figures 5, 11). Additionally, under the null hypothesis, some of the observed expression variance may be explained by increasing the estimated evolutionary variance, so power is reduced for genes with high *β*_*i*_.

**Figure 3:** The test for a gene with *β*_*i*_ varying from 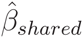 was computed for each gene. Those likelihood ratio test statistics (*LRT*_*β*_*i*_≠*β*_*shared*__) are plotted against the log of the *β* parameter estimated for each gene 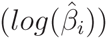 in a volcano plot. The dashed line indicates the value of 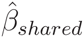.

We additionally estimated parametric bootstrapped *p*-values using 1000 simulations for each gene, finding that they roughly follow a uniform distribution with some excess of low *p*-values (Supplementary Figure 15), as is expected under our prediction that most genes are well described by *β*_*shared*_, while for a small number of genes *β*_*i*_ ≠ *β*_*shared*_. We compared those bootstrapped *p*-values to *LRT*_*β*_*i*_≠*β*_*shared*__ and found a clear correlation (Supplementary Figure 16). Using 1000 simulations, the minimum *p*-value is 0.001, so more simulations would be needed to more accurately assess the degree of departure from the null distribution in the tail of the distribution.

#### Candidate genes for expression adaptation and plasticity

Genes in the tail of the *LRT*_*β*_*i*_≠*β*_*shared*__ distribution with high 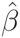 have conserved mean expression levels across species, but high variance within-species. A likely explanation is that the expression of these genes is highly plastic and that the genes are responding to individual environmental conditions. Among the most significant high 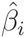 genes, we see PPIB, which has been implicated in immunosuppression (Price et al. 1991; Luban et al. 1993) and HSPA8, a heat shock protein (Daugaard et al. 2007) (Figure 4a). Based on their function, the expression levels of both of these genes are expected to vary depending on environmental inputs such as pathogen load and temperature. Since most of the samples were collected without standardized conditions, these environmental factors are likely to vary over individuals.

**Figure 4:** Each plot shows the expression profile across the 15 species for gene in the extreme tails of the empirical distribution of the test statistic for a gene-specific *β*_*i*_ differing from *β*_*shared*_ (*LRT*_*β*_*i*_≠*β*_*shared*__). (a) shows genes with high 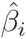 values and (b) shows genes with low 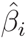 values.

Conversely, genes with low 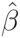 have unusually high evolutionary variance as compared to population variance, which is expected in cases of directional selection on expression level. The most extreme outlier with low 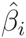 is F10, which encodes Factor X, a key blood coagulation protein produced in the liver (Uprichard and Perry 2002). F10 is highly expressed in armadillo as compared to the other mammals considered (Figure 4b). High F10 expression in armadillos may be caused by an environmental condition specific to armadillos, or by fixed genetic differences. We can not eliminate the possibility of an environmental factor underlying high F10 expression in armadillos without conducting experiments in controlled conditions. However, it has previously been found that armadillo blood coagulates two to five times faster than human blood (Lewis and Doyle 1964). A likely molecular cause is the increased expression of F10 observed here.

These results, together with the simulation results presented in the previous sections, suggest that the phylogenetic ANOVA application of the EVE model provides a versatile tool for identifying genes with relative elevated expression variance within-species, possibly due to plastic gene expression, or relative elevated expression divergence between-species, possibly due to species or lineage specific adaptive changes in gene expression. We emphasize that claims of adaptation would have to be followed up by additional lines of evidence.

### Testing for Branch-Specific Expression Level Shifts

The EVE model can be used to formulate hypotheses about branch-specific shifts in the expression of gene *i* by comparing likelihoods under 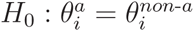 versus 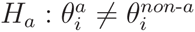, where 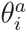 is the value of *θ*_*i*_ at all nodes in the shifted lineage(s), *a*, and 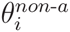 is the value of *θ*_*i*_ at the remaining (non-a) nodes. The corresponding likelihood ratio test statistic is asymptotically 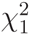 distributed. The phylogeny used for these analyses seems sufficient to achieve that asymptotic distribution for most genes (Supplementary Figure 17). We performed this test querying expression level shift on both the catarrhine (containing humans, chimpanzees, rhesus macaques, and vervets) and human lineages (Supplementary Tables 2, 3).

#### Candidate genes for adaptation on catarrhine and human lineages

In the test for expression shift in catarrhines (cat), we identify a number of interesting outliers (Supplementary Figure 18). The most significant shift is seen in DEXI, with higher expression level in catarrhines. This expression shift alone does not allow us to distinguish between environmental and genetic causation. However, studies in humans have shown high expression of DEXI to be protective against auto-immune diseases including type I diabetes and multiple sclerosis (Davison et al. 2012). If expression function is conserved across catarrhines, this suggests that increased DEXI expression in catarrhines may play an important role in immune response management.

Similarly, the test for expression shift on the human (hum) branch revealed interesting outliers (Supplementary Table 3), notably, two genes linked to fat metabolism or obesity. In the extreme tail of the distribution, we detected human-specific increased expression of MGAT1, which aids in metabolism of fatty acids to triglycerides (Yen et al. 2002), and the expression of which has been associated with excess retention of lipids (Lee et al. 2012). Additionally, we see that TBCA, a tubulin cofactor which assists in the folding of *β*-tubulin (Tian et al. 1996), has increased expression in humans. Given that reduced expression of TBCA through a heterozygous deletion has been associated with childhood obesity in humans (Glessner et al. 2010), it is possible that the human-specific increase in TBCA expression assists in metabolism of a high fat diet. However, in both cases, it is unclear if the increased expression in humans is an evolutionary shift in expression, helping to adapt to a diet more rich in fat, or if the increased expression in humans is environmentally responding to the diet. Expression level studies can only distinguish between these alternatives if the environmental conditions have been controlled between study objects, which for humans is only possible with cell line studies. Nonetheless, this new observation of human-specific regulatory changes for genes involved in fatty acid metabolism is interesting in light of the corresponding changes diet in humans.

Another gene with a significant expression shift in humans is BCKDK. BCKDK inactivates the branched-chain ketoacid dehydrogenase (BCKD) complex, which catalyzes metabolism of branched-chain amino acids (BCAAs). Nonsense and frame shift mutations in BCKDK have recently been linked to low levels of BCAAs and a phenotype including autism and epilepsy (Novarino et al. 2012). The observed increased human BCKDK expression may slow the metabolism of BCAAs so they can be processed into neurotransmitters (Novarino et al. 2012). Again, whether this shift has an adaptive genetic basis, or is a plastic response to human-specific conditions remains unclear.

#### Comparing results using the EVE model and species mean model

We compared our results for the expression shift tests to those reported in an analysis of the same data by Perry et al. (2012) using the species mean model described by Bedford and Hartl (2008). The distributions of 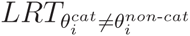 and 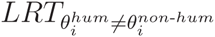 from that analysis deviate substantially from the 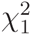 distribution expected under the null hypothesis (Supplementary Figure 20). This could be due to a number of possible numerical, optimization, or book-keeping errors, as these methods require a number of important technical considerations. In a comparison of the rank of expression shift test statistics as computed by Perry et al. (2012) and as computed using the EVE model, we see a general lack of correlation with some similarity in the extreme outliers discussed in that paper (Supplementary Figure 21).

To investigate if the results in Perry et al. (2012) were due to numerical problems we re-implemented the method and compared our results with those previously published by Perry et al. (2012). In our implementation, we see that the empirical distribution of test statistics are approximately 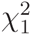 distributed with some excess of high values (Supplementary Figure 22) and a much improved correlation to EVE model test statistics (Figure 5), suggesting that the strong deviations for a 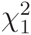 distribution in the Perry et al. (2012) results are largely due to numerical or optimization errors.

**Figure 5:** Each plot shows (a) 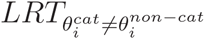 and (b) 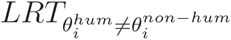 calculated using the EVE model (y-axes) and species mean model (x-axes) as implemented in this analysis. The line indicates *x* = *y*.

We then proceeded to compare the new results under the species mean model to the results of the EVE model. While both models identify similar genes with branch-specific *θ*_*i*_ shifts, we see much higher correlation between models for a shift on the catarrhine lineage than on the human lineage (Figure 5). Since the species mean model ignores variation within-species, it may identify genes where the mean expression appears to have shifted, even if the degree of variance may make that shift seem less extreme. By the same token, the EVE method may identify genes with a shift that cannot be explained by the expected within-species variance. This difference is most pronounced when considering shift of a single species (such as humans) where considering variance within that single species may alter the perception of an expression shift.

Figure 6 shows the three genes with the biggest difference in value of 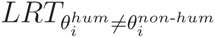 between the EVE and species mean models, that is, the genes that are most clearly identified by one model, while missed by the other. The gene TBCA, discussed above as a candidate for diet-associated expression adaptation, is a clear outlier under the EVE model 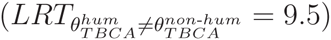, but is less easily identified using the species mean model 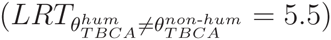. These results illustrate the importance of including within-species variance in the analyses of expression data evolution.

**Figure 6:** Each plot shows the expression profile for genes identified with an expression shift in humans by the EVE model, but not by the species mean (SM) model (top row), and identified by the species mean model, but not by the EVE model (bottom row). Expression levels in humans are highlighted in pink. Each plot shows 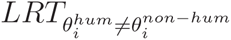 (as LRT) as computed under the EVE and species mean models.

## Discussion

We have described the EVE model for gene expression evolution which parameterizes the ratio between population and evolutionary variance in terms of a parameter *β* so that, in addition to more classic tests for selection on gene expression level, hypotheses regarding diversity to divergence ratios can be tested. We have explored a test for gene-specific *β*_*i*_, showing that the null distribution of the test statistic *LRT*_*βi ≠ βshared*_ is asymptotically 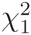, though depending on the size of the dataset and the value of the parameters, the null distribution may not have converged to the asymptote. We show that in these cases, a parametric bootstrap approach can be used to more accurately assess the significance of *LRT*_*βi ≠ βshared*_ values. Since the parametric bootstrap may be sensitive to variance in parameter estimates, it is prudent to verify its effectiveness on a particular data set with simulations before using it to interpret data.

The test for gene-specific *β*_*i*_ can be thought of as a phylogenetic ANOVA, or as a gene expression analog to the HKA test. This enables a previously unavailable line of inquiry into gene expression divergence, which may be indicative of expression-level adaptation to different environmental factors between species, and gene expression diversity, which may be indicative of plastic expression levels responding to environmental conditions. By utilizing a comparative approach, we can distinguish between genes which have high variance in expression levels within a species simply because expression of this gene has little effect on fitness, so is subject to drift, and genes with functional conserved expression levels across species along with high expression variance within-species because the gene mediates a plastic response to the environment. We have shown that by accounting for phylogeny our method has substantially improved power and reduced false positive rate as compared to traditional ANOVA, analogous to other results (Martins et al. 2002).

In applying the gene specific *β*_*i*_ test to a mammalian dataset, we identified several candidates for expression level divergence, most notably high expression of F10 in armadillos, which may be linked to their phenotype of rapid blood coagulation. We additionally identified several candidate genes for environmentally-responsive expression levels including PPIB, which helps regulate immunosuppression, and HSPA8, a heat shock protein. The identification of these biologically plausible candidates demonstrates the effectiveness of our method.

In addition to the novel test for unusual population or evolutionary variance, we used the EVE model to test for branch-specific shifts in expression level, as had been done previously with the species mean model (Hansen 1997; Butler and King 2004). Note that while the test for expression divergence may detect genes with branch-specific shifts, this more targeted test will detect shifts in expression on particular specified lineages. We found an increase in DEXI expression in catarrhines, which may have an adaptive role in auto-immune regulation to the catarrhine-specific pathogenic load. In humans, we found increased expression of two genes thought to be involved in lipid metabolism (MGAT1 and TBCA) and of BCKDK, the low expression of which has been linked to BCAA (necessary for neurotransmitters) deficiency, epilepsy, and autism.

When comparing our lineage-specific expression shift results to those previously reported using the species mean model, we observed startling differences. We attribute these differences primarily to a numerical or optimization problem in that original analysis, highlighting the importance of carefully addressing these issues. We performed an additional analysis using the species mean model to create a fair comparison. From that secondary analysis, we observe important differences between the EVE model and species mean model, most notably when testing for a shift in a single species. By discarding population variance, the species mean model may mistake a mild expression shift attributable to expected within-species variance for an evolutionary shift. We see this illustrated by the identification of an expression shift in humans for TBCA using the EVE model, but not using the species mean model.

As described here, the EVE model assumes one consistent and reliable phylogeny for all genes. Incomplete lineage sorting would violate this assumption, leading to unpredictable model behavior. To compensate, a Bayesian MCMC approach may be used to estimate the probability of expression data under a variety of underlying phylogenies using a method such as MrBayes (Ronquist and Huelsenbeck 2003). Additionally, like other similar tools, the EVE model and analyses described here do not account for expression correlations between genes, but rather, treat each gene independently. Gene expression data may be better described using a more complex multivariate approach (Dunn et al. 2013). Another important caveat is that while the EVE model is well-suited to detect adaptive divergence or plasticity of expression, this does not rule out increases in plasticity or canalization as part of the adaptive process (Lewontin 1974; Lande 1976).

The analyses described here provide examples of how the EVE model can be parameterized to test for expression divergence, diversity, or branch-specific shift. The tests for expression divergence and diversity can be used to identify genes with expression subject to different types of selection. For phylogenies where some species are known to be adapted to different environmental conditions, the branch-specific expression shift test can be formulated to identify genes with changes in expression that putatively underlie that adaptation. By changing parameter constraints, the EVE model can be used to test a variety of additional hypotheses. For example, tests may be formulated for branch-specific βvalues, which may be expected under branch-specific tightening or relaxation of constraint, or under unusual branch-specific demographic processes. The EVE model could also be used to test hypotheses of gene class-specific (rather than gene-specific) *β* values, which may vary based on gene class function. For example, genes involved in stress response may have a higher *β* value than housekeeping genes.

Like all comparative expression methods, the EVE method applies to any heritable quantitative trait with environmental components, including metabolomics (Nicholson and Lindon 2008; Cui et al. 2008; Sreekumar et al. 2009) and genome-wide methylation (Pokholok et al. 2005; Pomraning et al. 2009). As larger expression and other quantitative trait comparative datasets emerge, the versatile EVE model and framework described here will facilitate a wide variety of sophisticated analyses.

## Acknowledgments

We are immensely grateful to the individuals whose RNA samples were used in this study, without which none of this work would be possible. We thank George Perry and colleagues for making their data and results available and for assisting us in their interpretation, Youna Hu, Josh Schraiber, Tyler Linderoth, Julien Roux, Joe Felsenstein, Luke Harmon, Frank Anderson, and two anonymous reviewers for their valuable discussions on these topics, and Alex Safron for his help drawing schematics. This work was supported in part by National Institutes of Health grant 2R14003229-07, and National Science Foundation award 1103767. The funders had no role in study design, data collection and analysis, decision to publish, or preparation of the manuscript.

*

## References

Ackermann, R. R. and J. M. Cheverud. 2002. Discerning evolutionary processes in patterns of tamarin (genus saguinus) craniofacial variation. American Journal of Physical Anthropology Pages 260–271.

Beaulieu, J., D.-C. Jhwueng, C. Boettiger, and B. O’Meara. 2012. Modeling stabilizing selection: Expanding the Ornstein-Uhlenbeck model of adaptive evolution. Evolution 66:2369–2383.

Bedford, T. and D. Hartl. 2009. Optimization of gene expression by natural selection. Proc Natl Acad Scie USA 106:1133–1138.

Boettiger, C., G. Coop, and P. Ralph. 2012. Is your phylogeny informative? Measuring the power of comparative methods. Evolution 66:2240–2251.

Brawand, D., M. Soumillon, A. Necsulea, P. Julien, G. Csrdi, P. Harrigan, M. Weier, A. Liechti, A. Aximu-Petri, M. Kircher, F. Albert, U. Zeller, P. Khaitovich, F. Grtzner, S. Bergmann, R. Nielsen, S. Pääbo, and H. Kaessmann. 2011. The evolution of gene expression levels in mammalian organs. Nature 478:343–348.

Brent, R. 1973. Chapter 4. in Algorithms for minimization without derivatives (B. Dejon and P. Henrici, eds.). Prentice-Hall, Englewood Cliffs, NJ.

Broyden, C. 1970. The convergence of a class of double-rank minimization algorithms. Journal of the Institute of Mathematics and Its Applications 6:76–90.

Bullard, J., E. Purdom, K. Hansen, and S. Dudoit. 2010. Evaluation of statistical methods for normalization and differential expression in mRNA-Seq experiments. BMC Bioinformatics 11:94.

Butler, M. and A. King. 2004. Phylogenetic comparative analysis: A modeling approach for adaptive evolution. American Naturalist 164:683–695.

Butler, M., T. Schoener, and J. Losos. 2000. The relationship between sexual size dimorphism and habitat use in greater antillean anolis lizards. Evolution 54:259–272.

Cui, Q., I. A. Lewis, A. D. Hegeman, M. E. Anderson, J. Li, C. F. Schulte, W. M. Westler, H. R. Eghbalnia, M. R. Sussman, and J. L. Markley. 2008. Metabolite identification via the madison metabolomics consortium database. Nature Biotechnology 26:162–164.

Daugaard, M., M. Rohde, and M. Jäättelä. 2007. The heat shock protein 70 family: Highly homologous proteins with overlapping and distinct functions. FEBS Letters 581: 3702–676 3710.

Davison, L. J., C. Wallace, J. D. Cooper, N. F. Cope, N. K. Wilson, D. J. Smyth, J. M. Howson, N. Saleh, A. Al-Jeffery, K. L. Angus, H. E. Stevens, S. Nutland, S. Duley, R. M. Coulson, N. M. Walker, O. S. Burren, C. M. Rice, F. Cambien, T. Zeller, T. Munzel, K. Lackner, S. Blankenberg, P. Fraser, B. Gottgens, and J. A. Todd. 2012. Long-range DNA looping and gene expression analyses identify DEXI as an autoimmune disease candidate gene. Human Molecular Genetics 21:322–333.

Dunn, C. W., X. Luo, and Z. Wu. 2013. Phylogenetic analysis of gene expression. Integrative and Comparative Biology 53:847–856.

Felsenstein, J. 1985. Phylogenies and the comparative method. American Naturalist 125:1–15.

Felsenstein, J. 1988. Phylogenies and quantitative characters. Annual Review of Ecology and Systematics 19:445–471.

Felsenstein, J. 2008. Comparative methods with sampling error and within-species variation: Contrasts revisited and revised. The American Naturalist 171:713–725.

Fletcher, R. 1970. A new approach to variable metric algorithms. Computer Journal 13:317–322.

Fraser, H. 2011. Genome-wide approaches to the study of adaptive gene expression evolution. Bioessays 33:469–477.

Garamszegi, L., ed. 2014. Modern phylogenetic comparative methods and their application in evolutionary biology. 4 ed. Springer, New York.

Garland, T., A. Dickerman, C. Janis, and J. Jones. 1993. Phylogenetic analysis of covariance by computer simulation. Systematic Biology 42:265–292.

Gilad, Y., A. Oshlack, and S. Rifkin. 2006a. Natural selection on gene expression. Trends in Genetics 22.

Gilad, Y., A. Oshlack, G. Smyth, T. Speed, and K. White. 2006b. Expression profiling in primates reveals a rapid evolution of human transcription factors. Nature 440:242–245.

Glessner, J. T., J. P. Bradfield, K. Wang, N. Takahashi, H. Zhang, P. M. Sleiman, F. D. Mentch, C. E. Kim, C. Hou, K. A. Thomas, M. L. Garris, S. Deliard, E. C. Frackelton, F. G. Otieno, J. Zhao, R. M. Chiavacci, M. Li, J. D. Buxbaum, R. I. Berkowitz, H. Hakonarson, and S. F. Grant. 2010. A genome-wide study reveals copy number variants exclusive to childhood obesity cases. American Journal of Human Genetics 87:661–666.

Goldfarb, D. 1970. A family of variable metric updates derived by variational means. Mathematics of Computation 24:23–26.

Grafen, A. 1989. The phylogenetic regression. Philosophical Transactions of the Royal Society of London. Series B, Biological 326:119–157.

Gu, X. 2004. Statistical framework for phylogenomic analysis of gene family expression profiles. Genetics 167:531–542.

Hansen, T. 1997. Stabilizing selection and the comparative analysis of adaptation. Evolution 51:1341–1351.

Hansen, T. and K. Bartoszek. 2012. Interpreting the evolutionary regressions: The interplay between observational and biological errors in phylogenetic comparative studies. Systematic Biology 61:413–425.

Hansen, T., J. Pienaar, and S. Orzack. 2008. A comparative method for studying adaptation to a randomly evolving environment. Evolution 62:1965–1977.

Hansen, T. F. 2012. Adaptive landscapes and macroevolutionary dynamics. Pages 205–226 in The adaptive landscape in evolutionary biology (E. Svensson and R. Calsbeek, eds.). Oxford University Press.

Harmon, L. J. and J. B. Losos. 2005. The effect of intraspecific sample size on type I and type II error rates in comparative studies. Evolution 59:2705–2710.

Harmon, L. J., J. T. Weir, C. D. Brock, R. E. Glor, and W. Challenger. 2008. GEIGER: Investigating evolutionary radiations. Bioinformatics 24:129–131.

Housworth, E., E. Martins, and M. Lynch. 2004. The phylogenetic mixed model. The American Naturalist 163:84–96.

Hudson, R., M. Kreitman, and M. Aguadé. 1987. A test of neutral molecular evolution based on nucleotide data. Genetics 116:153–159.

Idaghdour, Y., W. Czika, K. Shianna, S. Lee, P. Visscher, H. Martin, K. Miclaus, S. Jadallah, D. Goldstein, R. Wolfinger, and G. Gibson. 2010. Geographical genomics of human leukocyte gene expression variation in southern morocco. Nature Genetics 42:62–67.

Ives, A., P. Midford, and T. Garland. 2007. Within-species variation and measurement error in phylogenetic comparative methods. Systematic Biology 56:252–270.

Kalinka, A., K. Varga, D. Gerrard, S. Preibisch, D. Corcoran, J. Jarrells, U. Ohler, C. Bergman, and P. Tomancak. 2010. Gene expression divergence recapitulates the developmental hourglass model. Nature 468:811–816.

Khaitovich, P., J. Kelso, H. Franz, J. Visagie, T. Giger, S. Joerchel, E. Petzold, R. E. Green, M. Lachmann, and S. Paäbo. 2006. Functionality of intergenic transcription: An evolutionary comparison. PLoS Genetics 2:e171.

Lande, R. 1976. Natural selection and random genetic drift in phenotypic evolution. Evolution 30:314–334.

Lande, R. 1979. Quantitative genetic analysis of multivariate evolution, applied to brain: Body size allometry. Evolution 33:402–416.

Lande, R. and S. J. Arnold. 1983. Measurement of selection on correlated characters. Evolution 37:1210–1226.

Larribe, F. and P. Fearnhead. 2011. On composite likelihoods in statistical genetics. Statistica Sinica 21:43–69.

Lee, Y., E. Ko, J. Kim, E. Kim, H. Lee, H. Choi, J. Yu, H. Kim, J. Seong, K. Kim, and J. Kim. 2012. Nuclear receptor PPAR*Γ*-regulated monoacylglycerol O-acyltransferase 1 (MGAT1) expression is responsible for the lipid accumulation in diet-induced hepatic steatosis. Proceedings of the National Academy of Sciences of the United States of America 109:13656–13661.

Lewis, J. H. and A. P. Doyle. 1964. Coagulation, protein and cellular studies on armadillo blood. Comparative Biochemistry and Physiology 12:61 – 66.

Lewontin, R. 1974. The analysis of variance and the analysis of causes. American Journal of Human Genetics 26:400–411.

Luban, J., K. L. Bossolt, E. K. Franke, G. V. Kalpana, and S. P. Goff. 1993. Human immunodeficiency virus type 1 Gag protein binds to cyclophilins A and B. Cell 73:1067 – 1078.

Lynch, M. 1991. Methods for the analysis of comparative data in evolutionary biology. Evolution 45:1065–1080.

Marroig, G. and J. M. Cheverud. 2004. Did natural selection or genetic drift produce the cranial diversification of neotropical monkeys? The American Naturalist 163:417–428.

Martins, E., J. Diniz-Filho, and E. Housworth. 2002. Adaptive constraints and the phylogenetic comparative method: A computer simulation test. Evolution 56:1–13.

Martins, E. and T. Hansen. 1997. Phylogenies and the comparative method: A general approach to incorporating phylogenetic information into the analysis of interspecific data. The American Naturalist 149:646–667.

Martins, E. P. and J. Lamont. 1998. Estimating ancestral states of a communicative display: a comparative study of cyclura rock iguanas. Animal Behaviour 55:1685–1706.

Necsulea, A., M. Soumillon, M. Warnefors, A. Liechti, T. Daish, U. Zeller, J. C. Baker, F. Grutzner, and H. Kaessmann. 2014. The evolution of lncRNA repertoires and expression patterns in tetrapods. Nature 505:635–640.

Nicholson, J. and J. Lindon. 2008. Systems biology: Metabolomics. Nature 455:1054–1065.

Nielsen, R. 2005. Molecular signatures of natural selection. Ann. Rev. Genet. 39:197–218.

Novarino, G., P. El-Fishawy, H. Kayserili, N. A. Meguid, E. M. Scott, J. Schroth, J. L. Silhavy, M. Kara, R. O. Khalil, T. Ben-Omran, A. G. Ercan-Sencicek, A. F. Hashish, S. J. Sanders, A. R. Gupta, H. S. Hashem, D. Matern, S. Gabriel, L. Sweetman, Y. Rahimi, R. A. Harris, M. W. State, and J. G. Gleeson. 2012. Mutations in BCKD-kinase lead to a potentially treatable form of autism with epilepsy. Science 338:394–397.

Nuzhdin, S., M. Wayne, K. Harmon, and L. McIntyre. 2004. Common pattern of evolution of gene expression level and protein sequence in *Drosophila*. Molecular Biology and Evolution 21:1308–1317.

Oakley, T., Z. Gu, E. Abouheif, N. Patel, and W. Li. 2005. Comparative methods for the analysis of gene-expression evolution: An example of using yeast functional genomic data. Mol. Biol. Evol. 22:40–50.

O’Meara, B. C., C. Ané, M. J. Sanderson, and P. C. Wainwright. 2006. Testing for different rates of continuous trait evolution using likelihood. Evolution 60:922–933.

Pagel, M. 1999. Inferring the historical patterns of biological evolution. Nature 401:887–884.

Perry, G., P. Melsted, J. Marioni, Y. Wang, R. Bainer, J. Pickrell, K. Michelini, S. Zehr, A. Yoder, M. Stephens, J. Pritchard, and Y. Gilad. 2012. Comparative RNA sequencing reveals substantial genetic variation in endagered primates. Genome Research 22:602–610.

Pickrell, J., J. Marioni, A. Pai, J. Degner, B. Engelhardt, E. Nkadori, J.-B. Veyrieras, M. Stephens, Y. Gilad, and J. Pritchard. 2010. Understanding mechanisms underlying human gene expression variation with RNA sequencing. Nature 464:768–772.

Pokholok, D. K., C. T. Harbison, S. Levine, M. Cole, N. M. Hannett, T. I. Lee, G. W. Bell, K. Walker, P. A. Rolfe, E. Herbolsheimer, J. Zeitlinger, F. Lewitter, D. K. Gifford, and R. A. Young. 2005. Genome-wide map of nucleosome acetylation and methylation in yeast. Cell 122:517–527.

Pomraning, K. R., K. M. Smith, and M. Freitag. 2009. Genome-wide high throughput analysis of DNA methylation in eukaryotes. Methods 47:142–150.

Price, E. R., L. D. Zydowsky, M. J. Jin, C. H. Baker, F. D. McKeon, and C. T. Walsh. 1991. Human cyclophilin B: A second cyclophilin gene encodes a peptidyl-prolyl isomerase with a signal sequence. Proceedings of the National Academy of Sciences 88:1903–1907.

Revell, L. J. 2012. phytools: An R package for phylogenetic comparative biology (and other things). Methods in Ecology and Evolution 3:217–223.

Revell, L. J., L. J. Harmon, and D. C. Collar. 2008. Phylogenetic signal, evolutionary process, and rate. Systematic Biology 57:591–601.

Rohlf, F. J. 2001. Comparative methods for the analysis of continuous variables: geometric interpretations. Evolution 55:2143–2160.

Rohlfs, R., P. Harrigan, and R. Nielsen. 2014. Modeling gene expression evolution with an extended Ornstein-Uhlenbeck process accounting for within-species variation. Molecular Biology and Evolution 31:201–211.

Ronquist, F. and J. P. Huelsenbeck. 2003. MrBayes 3: Bayesian phylogenetic inference under mixed models. Bioinformatics 19:1572–1574.

Schraiber, J., Y. Mostovoy, T. Hsu, and R. Brem. 2013. Inferring evolutionary histories of pathway regulation from transcriptional profiling data. PLoS Computational Biology 9:e1003255.

Shanno, D. 1970. Conditioning of quasi-newton methods for function minimization. Mathematics of Computation 24:647–656.

Sreekumar, A., L. M. Poisson, T. M. Rajendiran, A. P. Khan, Q. Cao, J. Yu, B. Laxman, R. Mehra, R. J. Lonigro, Y. Li, M. K. Nyati, A. Ahsan, S. Kalyana-Sundaram, B. Han, X. Cao, J. Byun, G. S. Omenn, D. Ghosh, S. Pennathur, D. C. Alexander, A. Berger, J. R. Shuster, J. T. Wei, S. Varambally, C. Beecher, and A. M. Chinnaiyan. 2009. Metabolomic profiles delineate potential role for sarcosine in prostate cancer progression. Nature 457:910–914.

Tian, G., Y. Huang, H. Rommelaere, J. Vandekerckhove, C. Ampe, and N. J. Cowan. 1996. Pathway leading to correctly folded *β*-tubulin. Cell 86:287 – 296.

Uprichard, J. and D. J. Perry. 2002. Factor X deficiency. Blood Reviews 16:97 – 110.

Whitehead, A. and D. Crawford. 2006. Variation within and among species in gene expression: Raw material for evolution. Molecular Ecology 15:1197–1211.

Yang, Z. and M. dos Reis. 2011. Statistical properties of the branch-site test of positive selection. Molecular Biology and Evolution 28:1217–1228.

Yen, C.-L. E., S. J. Stone, S. Cases, P. Zhou, and R. V. Farese. 2002. Identification of a gene encoding MGAT1, a monoacylglycerol acyltransferase. Proceedings of the National 845 Academy of Sciences 99:8512–8517

